# Expression profiling of the mature *C. elegans* nervous system by single-cell RNA-Sequencing

**DOI:** 10.1101/737577

**Authors:** Seth R Taylor, Gabriel Santpere, Molly Reilly, Lori Glenwinkel, Abigail Poff, Rebecca McWhirter, Chuan Xu, Alexis Weinreb, Manasa Basavaraju, Steven J Cook, Alec Barrett, Alexander Abrams, Berta Vidal, Cyril Cros, Ibnul Rafi, Nenad Sestan, Marc Hammarlund, Oliver Hobert, David M. Miller

## Abstract

A single neuron and its synapses define the fundamental structural motif of the brain but the underlying gene expression programs that specify individual neuron types are poorly understood. To address this question in a model organism, we have produced a gene expression profile of >90% of the individual neuron classes in the *C. elegans* nervous system, an ensemble of neurons for which both the anatomy and connectivity are uniquely defined at single cell resolution. We generated single cell transcriptomes for 52,412 neurons that resolve as clusters corresponding to 109 of the canonical 118 neuron classes in the mature hermaphrodite nervous system. Detailed analysis revealed molecular signatures that further subdivide identified classes into specific neuronal subtypes. Notably, neuropeptide-related genes are often differentially expressed between subtypes of the given neuron class which points to distinct functional characteristics. All of these data are publicly available at our website (http://www.cengen.org) and can be interrogated at the web application SCeNGEA (https://cengen.shinyapps.io/SCeNGEA). We expect that this gene expression catalog will spur the goal of delineating the underlying mechanisms that define the developmental lineage, detailed anatomy, synaptic connectivity and function of each type of *C. elegans* neuron.

## Introduction

Brain structure and function are defined by a fundamental motif comprised of an individual neuron and the signals it sends and receives. Because distinct genetic programs are likely to specify each neuron type, the goal of building a molecular model of the brain requires a gene expression map at single-cell resolution. Although new profiling methods have been successfully used to catalog diverse neuron types in a variety of organisms^1–5^, the complexity of these nervous systems and the correspondingly incomplete knowledge of their anatomy and wiring have hindered the goal of linking gene expression to each kind of neuron *in vivo*. To overcome this challenge, we have produced single cell RNA-Seq profiles for individual neurons in *C. elegans*, a model organism with a simple nervous system of well-defined structure and connectivity. The complete anatomy and wiring diagram of the *C. elegans* nervous system have been reconstructed from electron micrographs of serial sections^6,7^. This approach identified 118 anatomically distinct classes among the 302 neurons in the mature hermaphrodite nervous system. We established the *C. elegans* Neuronal Gene Expression Map & Network (CeNGEN) consortium^8^ to generate transcriptional profiles of each neuron class, thereby bridging the gap between *C. elegans* neuroanatomy and the genetic blueprint that defines it. We used fluorescence activated cell sorting (FACS) to isolate subgroups of neurons from L4 stage larvae for single cell sequencing. This approach generated profiles of 52,412 neurons, representing 109 of the 118 canonical neuron classes. Subtle differences in wiring and reporter gene expression within these canonical neuron classes predicted a substantially larger array of neuron types each with a unique molecular profile^9^. We have substantiated this idea by detecting distinct clusters of single cell profiles that correspond to subtypes of neurons within at least five neuron classes. Our data complement single cell profiles from *C. elegans* neurons at earlier developmental stages^10,11^ and open the door for a comprehensive effort to connect neuron-specific gene expression with the generation and function of the *C. elegans* nervous system.

## Methods

### Nematode Strains

The following strains were used in this study:

OH10689 (*otIs355 [rab-3(prom1)∷2xNLS-TagRFP]* IV)

EG1285 (*lin-15B&lin-15A(n765); oxIs12 [unc-47p∷GFP + lin-15(+)]* X)

NC3582 *(oxIs12 [unc-47p∷GFP + lin-15(+)]* X; *otIs355 [rab-3(prom1)∷2xNLS-TagRFP]* IV)

VM484 *(akIs3 [nmr-1p∷GFP + lin-15(+)]* V)

NC3572 *(akIs3 [nmr-1p∷GFP + lin-15(+)]* V; *otIs355 [rab-3(prom1)∷2xNLS-TagRFP]* IV)

OH9625 *(otIs292 [eat-4∷mCherry + rol-6(su1006)])*

OH11746 (*pha-1(e2123)* III; *otIs447 [unc-3p∷mCherry + pha-1(+)]* IV)

OH11157 (*pha-1(e2123)* III; *otIs393 [ift-20∷:NLS-TagRFP + pha-1(+)])*

OH13470 (*him-5(e1490)* V; *otIs354 [cho-1(fosmid)∷SL2∷YFP∷H2B])*

NC3579 (*otIs354 [cho-1(fosmid)∷SL2∷YFP∷H2B]*; *otIs355 [rab-3(prom1)∷2xNLS-TagRFP]*

IV)

NC3580 *(zdIs13 [tph-1∷GFP]* IV; *hpIs202 [ceh-10p∷GFP + lin-15(+)])*

CZ631 *(juIs14 [acr-2∷GFP + lin-15(+)]* IV)

RW10754 *(stIs10447 [ceh-34p∷HIS-24∷mCherry + unc-119(+)])*

NW1229 *(dpy-20(e1362)* IV; *evIs111 [F25B3.3∷GFP + dpy-20(+)])*

NC3583 *(stIs10447 [ceh-34p∷HIS-24∷mCherry + unc-119(+)]; evIs111 [F25B3.3∷GFP + dpy-20(+)])*

OH15430 *pha-1(e2123)* III*; otIs669 [NeuroPAL 15]*

OH16160 *pha-1(e2123)* III*; otEx7430 [nlp-56∷GFP + pha-1 (+)]; otIs669 [NeuroPAL 15]*

OH16161 *pha-1(e2123)* III*; otEx7431 [nlp-17∷GFP + pha-1 (+)]; otIs669 [NeuroPAL 15]*

### Preparation of larval strains and dissociation

Worms were grown on 8P nutrient agar 150 mm plates seeded with *E. coli* strain NA22. To obtain synchronized cultures of L4 worms, embryos obtained by hypochlorite treatment of adult hermaphrodites were allowed to hatch in M9 buffer overnight (16-23 hours) and then grown on NA22-seeded plates for 45-48 hours. The developmental age of each culture was determined by scoring vulval morphology (>75 worms)^12^. Single cell suspensions were obtained as described^13,14^, with some modifications. Worms were collected and separated from bacteria by washing twice with ice-cold M9 and centrifuging at 150 rcf for 2.5 minutes. Worms were transferred to a 1.6 mL centrifuge tube and pelleted at 16,000 rcf for 1 minute. 250 μL pellets of packed worms were treated with 500 μL of SDS-DTT solution (20 mM HEPES, 0.25% SDS, 200 mM DTT, 3% sucrose, pH 8.0) for 2-4 minutes. In initial experiments, we noted that SDS-DTT treatment for 2 minutes was sufficient to dissociate neurons from the head and tail, but longer times were required for effective dissociation of neurons in the mid-body and ventral nerve cord. The duration of SDS-DTT was therefore selected based on the cells targeted in each experiment. For example, NC3582, OH11746, and *juIs14* L4 larvae were treated for 4 minutes to ensure dissociation and release of ventral cord motor neurons. NC3579 and NC3580 L4 larvae were treated with SDS-DTT for 3 minutes. All other strains were incubated in SDS-DTT for 2 minutes. Following SDS-DTT treatment, worms were washed five times by diluting with 1 mL egg buffer and pelleting at 16,000 rcf for 30 seconds. Worms were then incubated in pronase (15 mg/mL, Sigma-Aldrich P8811) in egg buffer for 23 minutes. During the pronase incubation, the solution was triturated by pipetting through a P1000 pipette tip for four sets of 80 repetitions. The status of dissociation was monitored under a fluorescence dissecting microscope at 5-minute intervals. The pronase digestion was stopped by adding 750 μL L-15 media supplemented with 10% fetal bovine serum (L-15-10), and cells were pelleted by centrifuging at 530 rcf for 5 minutes at 4 C. The pellet was resuspended in L-15-10, and single-cells were separated from whole worms and debris by centrifuging at 100 rcf for 2 minutes at 4 C. The supernatant was then passed through a 35-micron filter into the collection tube. The pellet was resuspended a second time in L-15-10, spun at 100 rcf for 2 minutes at 4 C, and the resulting supernatant was added to the collection tube.

### FACS Analysis

Sorting was performed on a BD FACSAria™ III equipped with a 70-micron diameter nozzle. DAPI was added to the sample (final concentration of 1 μg/mL) to label dead and dying cells. Our general strategy used fluorescent reporter strains to target subgroups of neurons. For example, we used an *eat-4∷mCherry* reporter (OH9625) to target glutamatergic neurons, and an *ift-20∷NLS-TagRFP* reporter (OH11157) to label ciliated sensory neurons. We used an intersectional labeling strategy with a nuclear-localized pan-neural marker (*otIs355 [rab-3(prom1)∷2xNLS-TagRFP]* IV) to exclude cell fragments labeled with cytosolic GFP markers (NC3582). In other cases, we used an intersectional strategy to exclude non-neuronal cells. For example, *ceh-34p∷HIS-24∷mCherry* is expressed in pharyngeal muscles, pharyngeal neurons, and coelomocytes. To target pharyngeal neurons, we generated strain NC3583 by crossing *stIs10447 [ceh-34p∷HIS-24∷mCherry]* with the pan-neural GFP marker *evIs111* to isolate the cells that were positive for both mCherry and GFP. Non-fluorescent N2 standards and single-color controls (in the case of intersectional labeling approaches) were used to set gates to exclude auto-fluorescent cells and to compensate for bleed-through between fluorescent channels. In some cases, we set FACS gates to encompass a wide range of fluorescent intensities to ensure capture of targeted cell types. This less stringent approach may contribute to the presence of non-neuronal cells in our dataset (see Results). Cells were sorted under the “4-way Purity” mask. Sorted cells were collected into L-15-33 (L-15 medium containing 33% fetal bovine serum), concentrated by centrifugation at 500 rcf for 12 minutes at 4° C, and counted on a hemocytometer. Single-cell suspensions used for 10x Genomics single-cell sequencing ranged from 300-900 cells/μL.

### Single-cell RNA sequencing

Each sample (targeting 5,000 or 10,000 cells per sample) was processed for single cell 3' RNA sequencing utilizing the 10X Chromium system. Libraries were prepared using P/N 1000075, 1000073, and 120262 following the manufacturer's protocol. The libraries were sequenced using the Illumina NovaSeq 6000 with 150 bp paired end reads. Real-Time Analysis software (RTA, version 2.4.11; Illumina) was used for base calling and analysis was completed using 10X Genomics Cell Ranger software (v2.1.1 for *otIs355* samples, v2.2.0 for NC3582, and v3.0.2 for all others). Samples were processed with the 10x Genomics v2 Chemistry, except for samples from *juIs14* and NC3583, which were processed with v3 Chemistry. Detailed experimental information is found in Table S1.

### Downstream Processing

Reads were mapped to the *C. elegans* reference genome WBcel235 from Ensembl (build 96). The default barcode filtering algorithm in Cell Ranger can fail to capture cells in some conditions, especially with cells with variable sizes and RNA content^15^. Neurons, in particular, tend to have lower UMI counts than other cell types and can be missed by the default algorithm^11^. We therefore used the EmptyDrops method (with a threshold of 100 UMIs for determining empty droplets) from the R package DropletUtils^15^ to determine which droplets contained cells. This approach detected significantly more cells than the Cell Ranger method, and we were able to confidently annotate these additional cells as neurons. Our strategy also allowed us to determine the background RNA content of the empty droplets (i.e., droplets not containing cells) in each experiment, which we used for batch effect correction (see below).

Quality control metrics were calculated for each dataset using the scater package^16^ for R, using the percentage of UMIs from the mitochondrial genes *nduo-1, nduo-2, nduo-3, nduo-4, nduo-5, nduo-6, ctc-1, ctc-2, ctc-3, ndfl-4, atp-6*, and *ctb-1*. Droplets with greater than ten percent of UMIs coming from mitochondrial genes were removed. Datasets from individual experiments were merged using Seurat (v3)^17^, and downstream analysis was done using the Monocle (version 2.99.3)^18–20^ and Seurat (v3) packages.

### Dimensionality reduction and batch correction

Our strategy for covering the entire *C. elegans* nervous system involved the independent isolation of multiple partially overlapping subgroups of neurons for single cell sequencing (**Figure 1C**, Table S1). During the downstream annotation of clusters, we detected multiple clusters corresponding to the same neuron that clustered separately by experiment (i.e., each cluster was comprised almost entirely of cells from only one experiment). One likely source of these batch effects is differing compositions of background RNA from lysed or damaged cells between experiments^11,21^. This background RNA should be present in the droplets with fewer than 50 UMIs (i.e., empty droplets that do not contain cells). We generated expression profiles of the empty droplets for each experiment. We then compared these profiles to the genes differentially expressed between clusters of the same neuron type that separated by experiment, and found that the background profile genes were highly differentially expressed in cells of the same type between experiments, indicating the background RNA was a major source of batch effects. We compiled a consensus list of 339 genes (Table S2) that were the most abundant in the empty drop RNA profiles across all the experiments. The downstream analysis steps of dimensionality reduction require the selection of highly variable genes. We used the Monocle analysis pipeline and estimated mean expression and dispersion parameters for all genes in the dataset. We selected genes to use for principle component analysis based on mean expression (> 0.002) and dispersion (>5) parameters. These parameter thresholds selected some of the genes in the empty drop RNA profile. To correct for batch effects, we manually removed these genes from use in principal component analysis (PCA) and other downstream analyses. This approach does not alter the underlying gene expression data. We then reduced the dimensionality of the dataset with PCA (125 principal components), followed by the Uniform Manifold Approximation and Projection (UMAP)^22,23^ algorithm in Monocle (reduceDimension function, parameters were default other than: min_dist = 0.2, n_neighbors = 75). We then clustered the cells using the Louvain algorithm in Monocle (res = 1e−3, k = 18).

**Figure 1.**
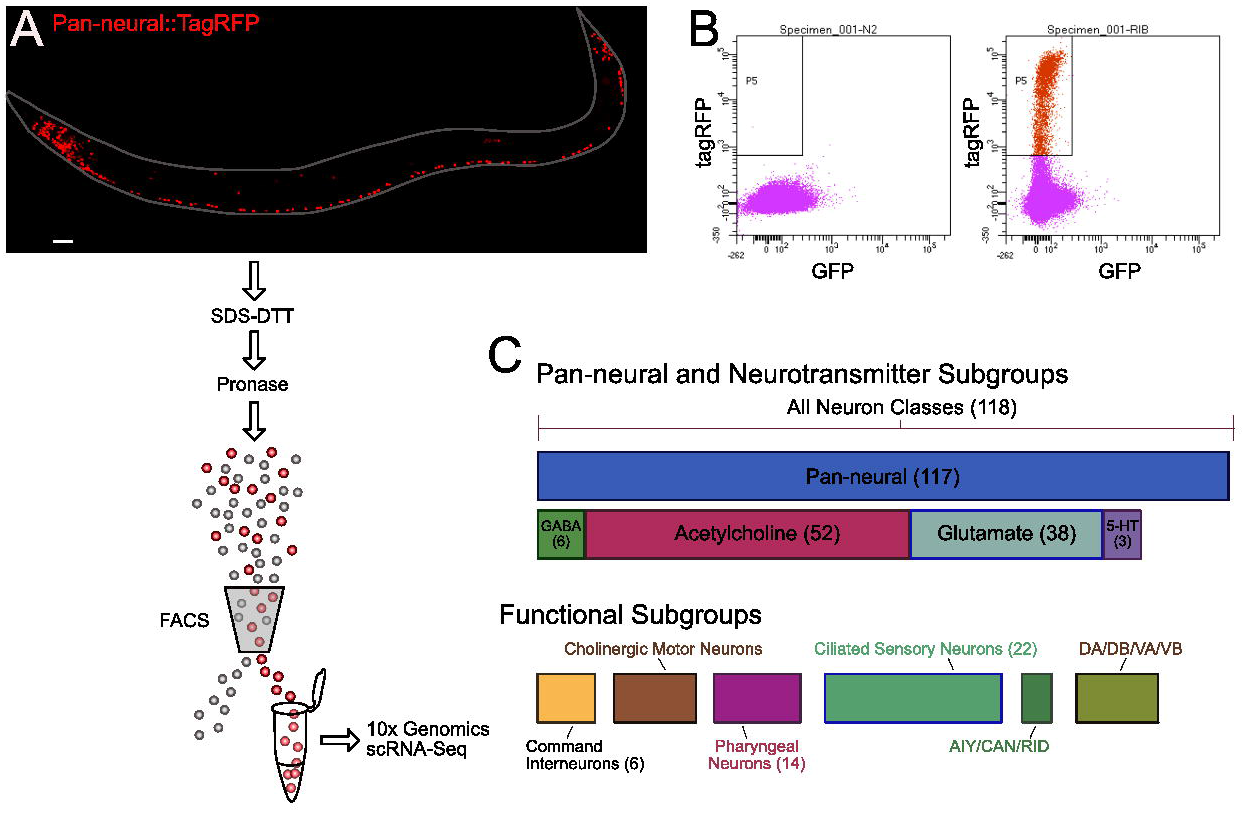
Overview of experimental procedure and labeling strategies. (**A**) Confocal image of pan-neural strain *otIs355* expressing nuclear-localized TagRFP in all neurons except CAN. L4 animals were treated with SDS-DTT and dissociated with pronase to produce single-cell suspensions. Targeted subgroups of neurons were isolated by Fluorescence Activated Cell Sorting (FACS) and collected for single-cell RNA sequencing using the 10x Genomics 3’ platform. B) Scatterplots of control N2 worms showing background autofluorescence (left panel) and *otIs355* worms showing TagRFP (red) labeled cells (right panel). Cells with a wide range of RFP expression above background were collected (the rectangle labeled P5) for single-cell sequencing. C) Graphical representation of the subgroups of neurons that were targeted in individual experiments. An initial pan-neural experiment was followed by experiments targeting cell types based on neurotransmitter type. Subsequent experiments targeted subgroups of neurons based on functional properties or specific patterns of gene expression to capture additional neuron types.

We assigned tissue and cell identity to the majority of cells in our dataset by relying on gene expression assignments from WormBase and recent literature (see Table S3). We manually excluded clusters we identified as doublets due to co-expression of cell type specific markers. Some clusters in the initial global dataset appeared to contain multiple closely related neuron types (i.e. dopaminergic neurons ADE and PDE, oxygen sensing neurons AQR, PQR, URX, touch receptor neurons ALM, AVM, PLM, AVM and pharyngeal neurons). Additional analysis of these separate clusters (i.e., reapplication of PCA, UMAP, and Louvain clustering to just these clusters) effectively separated many of these cell types into individual clusters.

### Gene expression analyses

Gene expression profiles for each neuron class were generated as described^10^. Linear regression was performed using the lm function in R. Expression data for a subset of genes are distorted by overexpression from co-selectable markers (*lin-15A, lin-15B, pha-1, rol-6, unc-119, dpy-20*) or from a gene-specific 3’ UTR included in fluorescent reporter constructs (*eat-4*, *unc-54*). These genes are annotated in the SCeNGEA web application.

For visualization of gene expression data in the web application and for differential gene expression tests, the data were imported into Seurat (v3) and the raw counts were normalized using the variance stabilizing transformation (VST) implemented in the function SCTransform with default parameters^17^. Differential gene expression tests used the Seurat v3 default Wilcoxon rank sum test.

### Reporter Strain Generation

GRP reporters for the neuropeptide genes *nlp-17* and *nlp-*56 were created by PCR Fusion^24^ whereby the 5’ intergenic region of the gene of interest and the coding sequence of GFP with 3’ UTR of *unc-54* were fused in subsequent PCR reactions. We used the entire intergenic region of the genes of interest, 2954bp for nlp-56 (forward primer: ggttcactggaataaatatatgcactgtatc, reverse primer: ctggaagagttgaatcatatggtttagaag) and 372bp for nlp-17 (forward primer: tcatctaaaatatattttcaaaacgattttctgtgc, reverse primer: attttctgtgaaaaagcctgactttttc) to make our reporters. The reporters were injected directly into NeuroPAL *pha-1* strain (OH15430 *pha-1(e2123); otIs669[NeuroPAL 15])*^25^ as a complex array with OP50 DNA (linearized with *sca-1)* and *pBX [pha-1 (+)]*^26^ as a co-injection marker. For *nlp-17*, the reporter, *pBX [pha-1 (+)]* and OP50 DNA were injected at a concentration of 10ng/ul, 6.2ng/ul and 99.96ng/ul, respectively. In the case of *nlp-56*, the reporter, *pBX [pha-1 (+)]* and OP50 DNA were injected at a concentration of 9.5ng/ul, 5.2ng/ul and 94.9ng/ul, respectively. After injection, animals were kept at 25°C for selection of the array positive worms and maintained for at least three generations before images were taken with a confocal microscope (Zeiss LSM880 confocal). The images were analyzed using Fiji Imaging Software to identify the neurons expressing the reporters.

## Results

Because adult neurons were inefficiently dissociated by our methods (data not shown), we isolated neurons from L4 larvae, since all neuron types are available by this developmental stage. In our initial attempt to profile the entire mature *C. elegans* hermaphrodite nervous system, we used FACS to isolate neurons from a pan-neural marker strain (*otIs355*) which expresses nuclear-localized TagRFP in all neuron classes except CAN^27^ (**Figure 1A-B**). Although this approach yielded a variety of distinct neuron types, selected classes were either underrepresented or absent (Table S4). To overcome this limitation, we isolated cells from a series of fluorescent marker strains that labeled distinct subsets of neuron types (**Figure 1C**, Table S1). This approach produced a combined dataset of 65,450 single cell transcriptomes. Application of the Uniform Manifold Approximation and Projection (UMAP)^22,23^ dimensional reduction algorithm effectively segregated most of these cells into distinct groups in the UMAP space (**Figure 2**). We clustered cells in the UMAP using the Louvain algorithm.^28^

**Figure 2.**
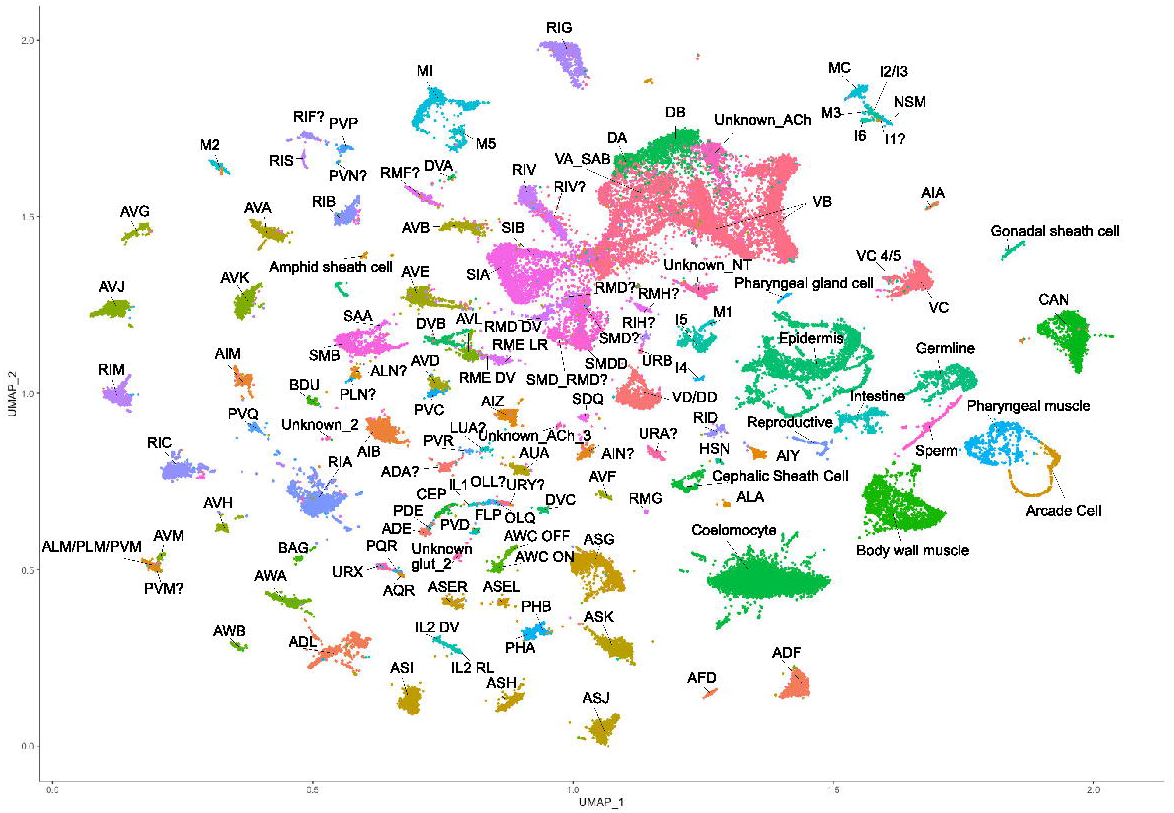
Global UMAP of all cells. UMAP projection of single cell profiles from the 65,450 cells that passed quality control. Cell type assignments are based on gene expression data from WormBase and from the literature. Annotated clusters corresponding to specific cell types are labeled (e.g., ADF) and distinguished by color. Non-neuronal cell types include coelomocytes, muscle cells, intestinal cells, reproductive system cells, and others.

We used expression data from WormBase, the literature, and unpublished studies from the Hobert lab to assign cell type identities to individual clusters (see Table S3 for genes used for annotation). We annotated 52,412 cells (79% of all single cell profiles) as neurons and detected several non-neuronal cell types. Among these were coelomocytes, body muscle cells, epidermal cells, glial cells, germline cells, intestinal cells, reproductive system cells, and non-neuronal pharyngeal cells. In some instances, these non-neuronal cells were labeled in the reporter strains. For example, the *zdIs13 [tph-1∷GFP]* transgene in NC3580 is expressed in non-neuronal pharyngeal cells. The wide gates used for the FACS isolation (designed to capture as many types of fluorescent cells as possible) could also account for the inclusion of weakly labeled non-neuronal cells. We noted that coelomocytes were most abundant in experiments using strains expressing mCherry (*otIs292* and *otIs447)*. This effect likely results from neurons shedding mCherry+ exophers, which are then taken up by coelomocytes^29^, causing them to be isolated along with mCherry-labeled neurons.

We created a separate neuronal UMAP (**Figure 3**) by filtering non-neuronal cells from the global dataset. We detected an average of 1461 UMIs and 460 genes per individual profiled neuron. The UMAP projection of the 52,412 neurons resolved many individual neuronal classes. Some clusters appeared to contain multiple closely related neuronal classes (i.e., dopamine neurons, touch receptor neurons, oxygen-sensing neurons, ventral cord motor neurons). Individual UMAP projections of these clusters allowed for more precise annotation (**Figure 4**). All together, we have confidently identified 93 of the 118 (78.8%) neuron classes in the mature hermaphrodite nervous system, and putatively identified another 16 (13.6%) neuron classes. The putative annotations represent our best guess based on marker expression and include question marks in the annotation (e.g. ADA?, AIN?). Overall, we successfully annotated 96.9% of the cells in the neuronal dataset.

**Figure 3.**
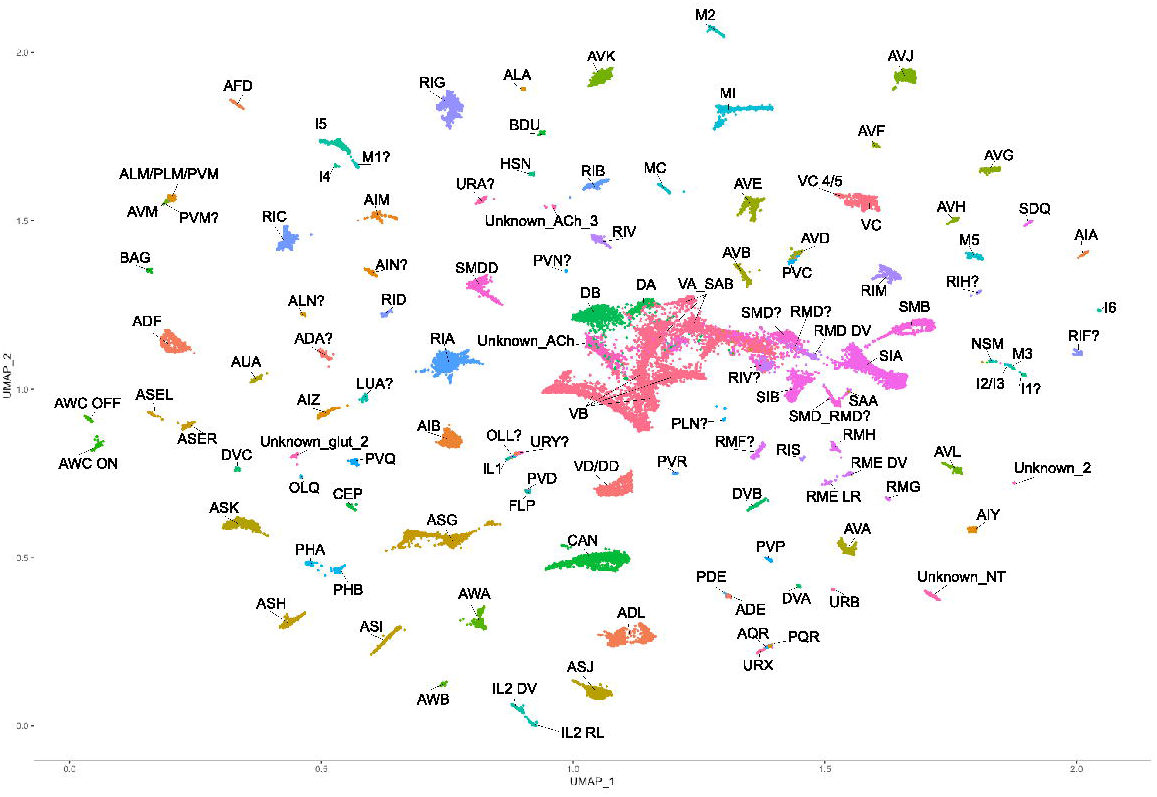
Neuronal UMAP. UMAP projection of 52,412 neurons after removal of non-neuronal cell types. Colors correspond to cell type. Most clusters have been assigned to a specific neuron type, and 4 of the 5 unidentified clusters are likely neuronal (the cluster labeled “Unknown_NT” may not be neuronal). Several neuronal classes segregate into distinct subclasses, including the sensory neuron pairs ASE and AWC, the inner labial neurons IL2, and the RME motor neurons. Some closely related cell types with sparse representation in the data cluster with related cell types (ADE/PDE, AQR/PQR/URX, ALM/AVM/PLM/PVM, and pharyngeal neurons). Neuronal assignments for these clusters were derived from separate sub-UMAPs (Figure 4).

**Figure 4.**
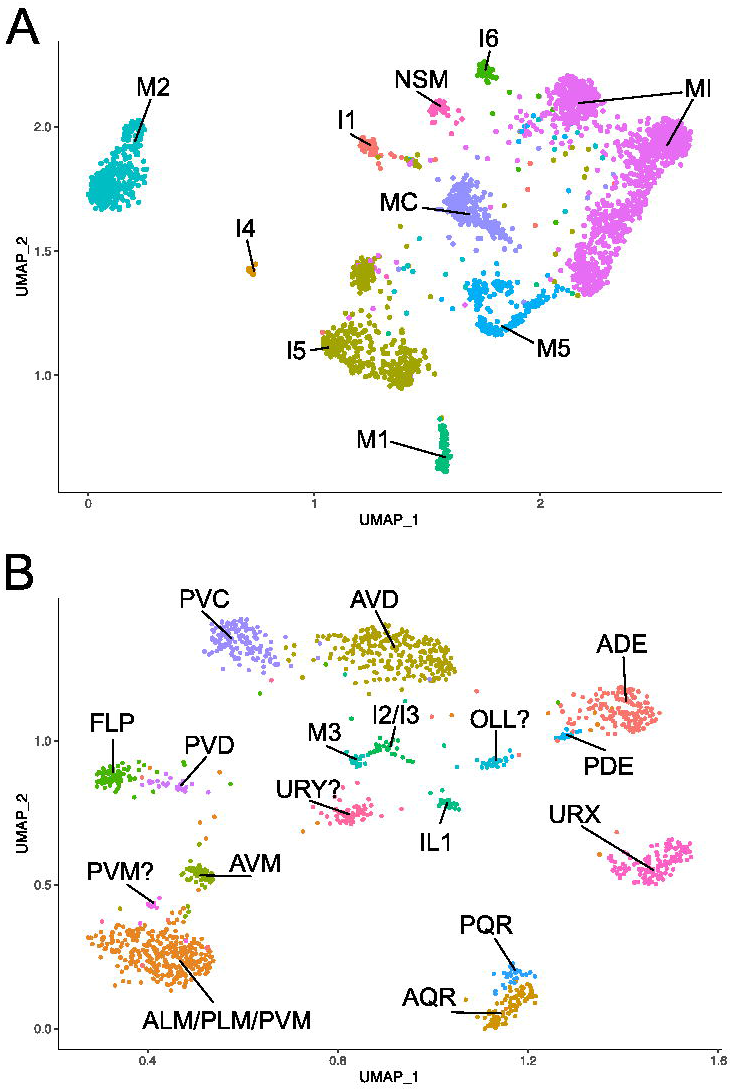
Sub-UMAPs resolve additional neuronal classes. A) UMAP projection of pharyngeal classes. Pharyngeal neurons cluster separately in the analysis, allowing for more precise annotation than in the neuronal UMAP projection (Figure 3). B) UMAP of neuron classes that clustered together in Figure 3. Application of PCA, UMAP and clustering on this subset of neurons resolved ADE, PDE, URX, AQR, PQR, FLP, PVD, AVD, PVC, AVM, IL1, OLL, URY, and possibly PVM. ALM/PLM are likely in the same cluster, and the pharyngeal neurons I2/I3 cluster together, which could be due to the small number of cells from these neuron classes in our dataset.

During our effort to assign neuron identities to each cluster, we detected two neuropeptide genes (*nlp-17* and *nlp-56)* which were highly enriched in single unannotated clusters (**Figure 5A, C**). To determine the neuron types expressing these peptides, we generated transgenic GFP reporter strains consisting of the entire intergenic regions upstream of the predicted *nlp-17* and *nlp-56* start sites. To facilitate neuron identification, the reporter strains were built in the NeuroPAL background, a strain which unambiguously resolves all neuron types^25^. The *nlp-56∷GFP* reporter was only expressed in the RMG neurons in the head (**Figure 5B**). The *nlp-17∷GFP* reporter was only expressed in one pair of neurons in the tail, identified as PVQ (**Figure 5D)**. These clusters displayed additional markers for RMG and PVQ, respectively (Table S3), allowing for confident annotation of these neuron types. The specificity of these reporters highlights how these data can reveal cell-type specific expression patterns that enable genetic access to a wider range of neuron classes than has been previously possible.

**Figure 5.**
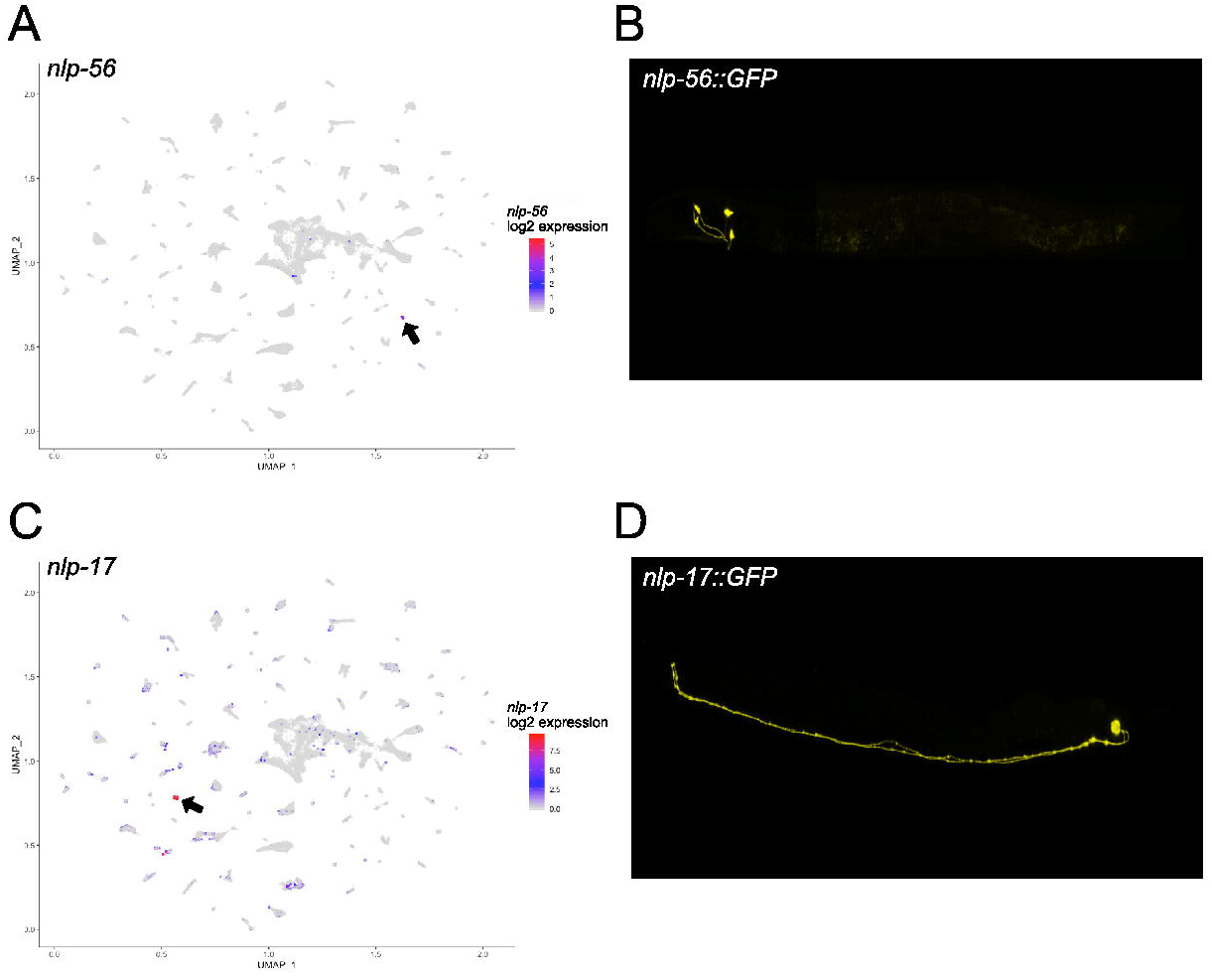
Neuron-type specific expression of *nlp* genes. A) Neuronal UMAP showing expression of *nlp-56.* Expression is limited almost entirely to one cluster (black arrow). B) Confocal image of an *nlp-56∷GFP* reporter strain at the L4 larval stage. The reporter is only expressed in one pair of neurons in the head identified as RMG using the NeuroPAL system. C) Neuronal UMAP displaying expression of *nlp-17.* The black arrow indicates the single cluster with high levels of *nlp-17*. A few other clusters show low levels of *nlp-17* expression. D) Confocal image of an L4 *nlp-17∷GFP* reporter. This reporter is expressed in a single pair of tail neurons, identified with NeuroPAL as PVQ.

Nine neuron classes (AS, M4, PDA, PDB, PHC, RIP, RIR, PVT, and PVW) have not been identified. Some of these classes may be represented in our dataset, as four distinct clusters appear to be neuronal (i.e., express neuronal markers *sbt-1* and *egl-21*), but have not been assigned to a specific neuron type. Because few marker genes have been previously annotated for several of these missing classes (e.g., PDB, RIR, RIP) and for many of the neuron types which have been “putatively” identified (e.g., PVN, RMF), we are using cluster-specific markers to produce new fluorescent reporters and single molecule fluorescence *in situ* hybridization (smFISH) probes for *ab initio* assignments of these clusters to specific neuron types. Some neuronal classes are likely embedded in clusters with closely related neuron types. These include the GABAergic VD and DD motor neurons which are in the same cluster, and the sparsely represented pharyngeal neurons I2 and I3 which appear in a single distinct cluster (**Figure 4B**). Touch sensitive neurons are grouped in three clusters; a large cluster likely containing ALM and PLM, a smaller cluster corresponding to AVM, and a small group of cells possibly corresponding to PVM. The motor neuron class SAB may be combined with the VA neurons.

Existing gene expression data indicate that many of the initially described 118 neuron classes^6,7^ can be subdivided into molecularly distinct subclasses^9^. For example, we detect individual clusters for two types of the bilateral asymmetric neuron pairs, ASE (ASER and ASEL) and AWC (AWC^ON^ and AWC^OFF^) as previously confirmed by single-cell embryonic and early larval RNA-seq datasets^10,11^. Additional distinctions resolved by our approach include dorsal/ventral vs left/right subtypes for inner labial class IL2 neurons, dorsal/ventral vs left/right subtypes for the GABAergic RME motor neurons and two distinct VC motor neuron clusters corresponding to VC4/VC5 vs all other VCs. We observed additional possible subclasses of the SIA, SMD, RMD, VA, and VB neurons that await further validation. Additional classes with 3, 4, or 6-fold symmetry (e.g., SAB, CEP, OLQ, SAA, URY, IL1) may also have molecularly distinct subclasses^9^. The low number of cells (< 75, in most cases) assigned to these classes could account for the absence of distinct subclasses.

To characterize the molecular differences between neuronal subclasses, we tested differential gene expression between the following subclasses: ASER v ASEL, AWC^ON^ v AWC^OFF^, IL2 DV v IL2 LR, RME DV v RME LR, and VC 4/5 v VC (Tables S5-S9). We required a gene to be detected in at least 10% of the cells in the higher-expressing cluster and used an adjusted p-value cut-off of 0.05. Strikingly, this analysis revealed that neuropeptides were often highly specific within neuron subclasses. We detected 105 genes differentially expressed between ASER and ASEL (Table S5). As expected, many genes of the guanylyl cyclase (*gcy)* family were differentially expressed^30–32^, including *gcy-14, gcy-17, gcy-6, gcy-7*, and *gcy-20* in ASEL, and *gcy-3*, *gcy-22, gcy-19* in ASER. The neuropeptides *ins-3, nlp-7, nlp-5, nlp-8, nlp-21* and *flp-9* were enriched in ASER compared to ASEL. We detected 58 genes differentially expressed between AWC^ON^ and AWC^OFF^ (Table S6). Several genes of the serpentine receptor class showed highly asymmetric expression, as previously reported^33–35^. Of 10 differentially expressed serpentine receptor GPCRs*, srd-5, srsx-3, srsx-5, srx-2, str-144*, and *str-116* were enriched in AWC^OFF^. Four were enriched in AWC^ON^ (*srt-24, srt-28, srt-42*, and *srx-50*). Neuropeptides differentially expressed between AWC^ON^ and AWC^OFF^ included *ntc-1, nlp-5, nlp-7*, and *flp-21*, which were enriched in AWC^ON^. 48 genes were differentially expressed between IL2 DV and IL2 LR (Table S7). In addition to neuropeptides (*flp-14* and *flp-9* were enriched in IL2 LR), calcium ion binding proteins and ion channels (*egas-1* in IL2 DV, *egas-4* in IL2 LR) distinguished the subclasses. We detected 38 genes differentially expressed between the left/right and dorsal/ventral GABAergic RME neurons (Table S8), including the neuropeptides *nlp-41, flp-16*, and *nlp-11*, which were enriched in RME DV compared to RME LR. We detected 81 genes differentially expressed between VC 4/5 and the other VC neurons (Table S9). Several neuropeptides were among the most significantly differentially expressed genes, including *flp-27, flp-24, nlp-7*, and *flp-22*, all of which were enriched in VC 4/5. The peptides *nlp-6, flp-9, nlp-13*, *flp-6, flp-2* and *flp-3* were highly enriched in the other VC neurons.

This specificity of expression led us to investigate the distribution of neuropeptides throughout the entire neuronal dataset (Table S10). Most of the neuropeptide genes were detected in the dataset. All neuron classes expressed at least one neuropeptide gene, and most expressed several peptides. Several neuropeptides were widely expressed in many neuron types, including *flp-9, flp-5*, and *nlp-21.* We also detected several peptides with expression restricted to just one or two neuron types, including exclusive expression of *flp-23* in HSN, *nlp-*2 in AWA, *flp-33* in ADE and PDE, *nlp-19* in AVH and putative ADA, and *nlp-20* in ALN, SMB, and putative PLN. These data also reveal that several closely related types of neurons display divergent neuropeptide profiles. The oxygen sensing neurons AQR, PQR and URX are quite similar transcriptionally (they cluster very closely together in the neuronal UMAP), These neuron types, however, have distinct profiles of neuropeptides. For example, among these three classes, *nlp-43* is only in AQR, *flp-4* is only expressed in PQR, and *flp-8* is only in URX (**Figure 6**). Overall, these data indicate that neuropeptide signaling is highly divergent between neuron types as well as within subclasses of the same neuron type, and that neuropeptide markers may be useful for further investigating neuronal diversity.

**Figure 6.**
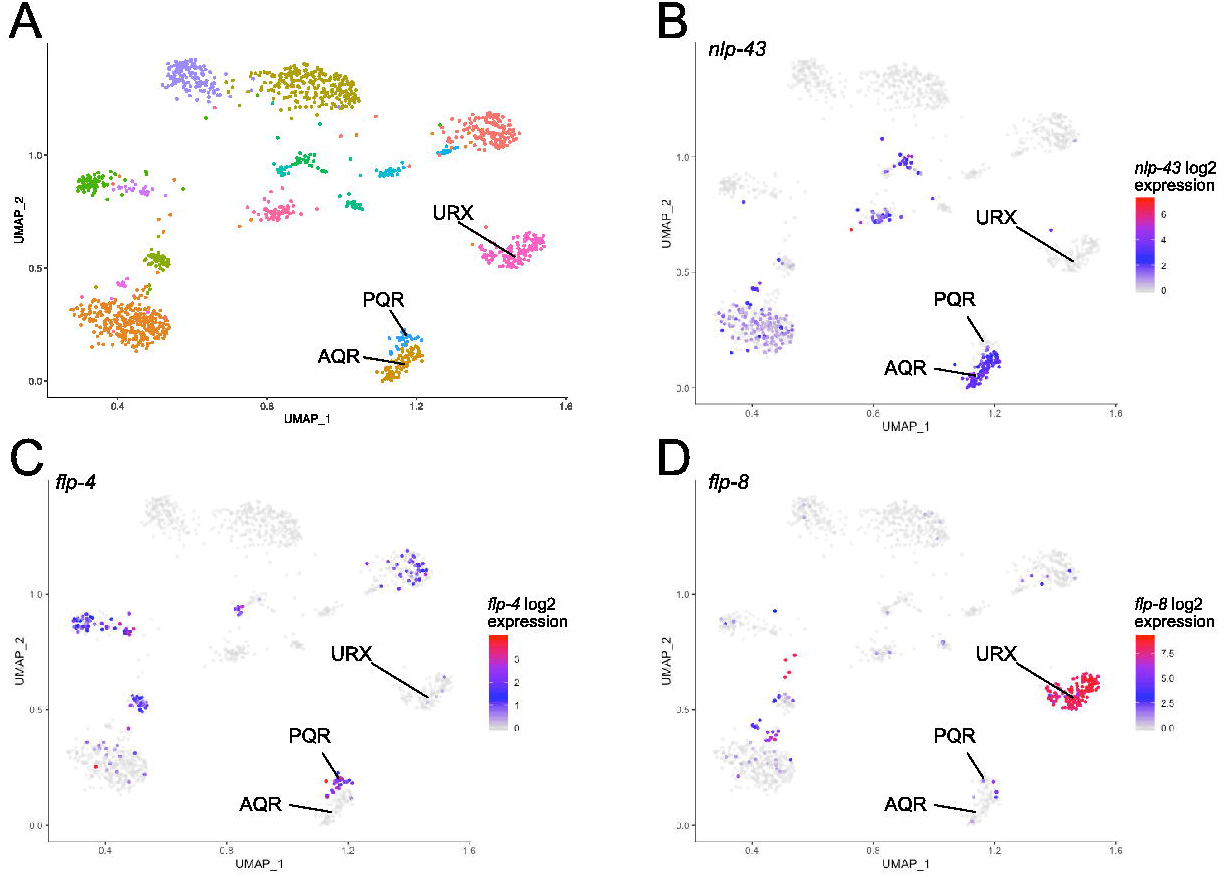
Neuropeptide expression differentiates closely-related neuron types. A) Sub-UMAP as in Figure 4 showing clusters corresponding to the oxygen sensing neurons AQR, PQR, and URX. B) Expression of *nlp-43* in AQR but not PQR or URX. C) Expression of *flp-4* in PQR but not AQR or URX. D) *flp-8* is expressed strongly in URX, but weakly in AQR and PQR.

The number of cells assigned to each neuron class ranged from 11 cells (putative PVN?) to 6706 cells (VB). The large number of VB cells arises primarily from one experiment using the *juIs14 [acr-2∷GFP]* strain, which labels DA, DB, VA, and VB motor neurons. We generated gene expression profiles for each neuron class to explore the relationship between the number of cells and the number of genes detected (defined as at least 1 UMI in 1 cell). The detection of transcripts in droplet-based single-cell RNA sequencing is a stochastic event, and the 10x Genomics Single Cell 3’ assay captures only a fraction of the mRNA in each cell^36^. Thus, a greater proportion of the transcriptome for a given cell type should be captured by sequencing more single cells from that population. A plot of the number of genes detected in each identified cluster vs the number of cells in that cluster confirms that clusters with larger numbers of cells also contain a greater number of overall genes (**Figure 7**). Linear regression analysis using a logarithmic function yielded an empirical formula to model this relationship: Number of genes = −4036 + 2101*ln(number of cells) (adjusted R-squared = 0.9181, p < 2.2 e-16). For example, within the range of cell numbers observed (i.e., 11 to 6706), the model predicts that a 10% increase in the number of cells for a given cluster will yield an additional 200 genes. Hence, the number of single cell profiles for a given neuron type must be considered in interpreting these gene expression data. In particular, genes that are actually expressed in a given cell type may fail to be detected (e.g., ‘dropouts’) in clusters with limited numbers of cells. Thus, our detected average of 7241 genes per neuron class (median of 7474 genes) is likely an underestimate of the total number of genes in a given neuron type *in vivo*.

**Figure 7.**
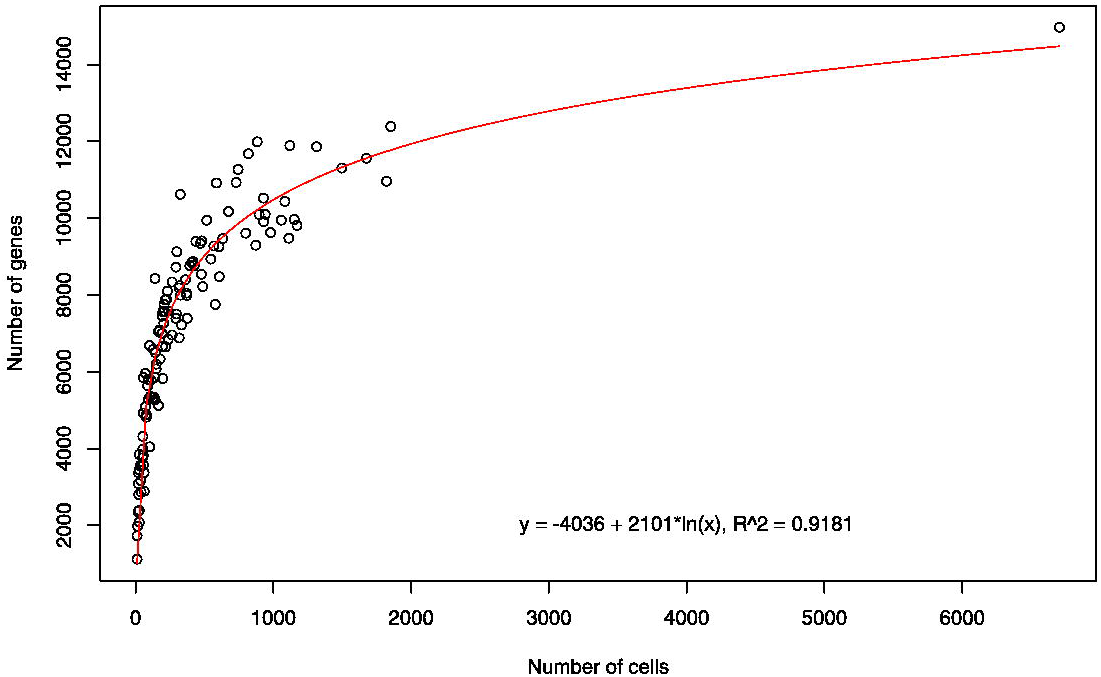
The number of genes detected for a cell type increases non-linearly with increasing numbers of cells. The number of genes detected (1 UMI in at least 1 cell) for each neuron class (range: 1116 to 14,970, median 7472) was plotted against the number of cells for each class (range: 11 to 6,706, median 228). Linear regression of a natural logarithmic function yielded the indicated best fit, with an adjusted R-squared value of 0.9181. This model predicts that a 10% increase in the number of cells will result in an additional 200 detected genes within the range of cells used to generate this plot.

CeNGEN will address this disparity with the parallel strategies of collecting additional single cell sequence data as well as generating deep bulk RNA-sequence data from FACS-isolated population of specific neurons types^8^. Whereas 10x Genomics single cell RNA sequencing provides expression data based on 3’ counting of polyA-mRNAs, bulk RNA-sequencing offers the additional benefits of whole transcriptome sequences and detection of non-coding RNAs, which are not captured by single cell sequencing. In ongoing work, we are analyzing these single cell data at a nervous system-wide level to elucidate gene expression patterns that underlie the development, connectivity, and function of the *C. elegans* nervous system.

We have developed a web application at http://cengen.shinyapps.io/SCeNGEA to explore the single-cell data generated by this project. Users can query for gene expression by neuron type, expression patterns for a given gene, and for differential gene expression between neuron types. The raw data have been deposited with the Gene Expression Omnibus (www.ncbi.nlm.nih.gov/geo) (in progress). The data as an R object and additional supporting files can be downloaded from the CeNGEN website, www.cengen.org.

## Acknowledgements

This work was funded by the NIH grant R01NS100547 to MH, OH, DMM, and NS, and by Vanderbilt Trans-Institutional Program funds to DMM. Flow Cytometry experiments were performed in the VMC Flow Cytometry Shared Resource. The VMC Flow Cytometry Shared Resource is supported by the Vanderbilt Ingram Cancer Center (P30 CA68485) and the Vanderbilt Digestive Disease Research Center (DK058404). The Vanderbilt VANTAGE Core provided technical assistance for this work. VANTAGE is supported in part by CTSA Grant (5UL1 RR024975-03), the Vanderbilt Ingram Cancer Center (P30 CA68485), the Vanderbilt Vision Center (P30 EY08126), and NIH/NCRR (G20 RR030956). Some strains were provided by the CGC, which is funded by NIH Office of Research Infrastructure Programs (P40 OD010440).

